# Structure of *Anabaena flos-aquae* gas vesicles revealed by cryo-ET

**DOI:** 10.1101/2022.06.21.496981

**Authors:** Przemysław Dutka, Lauren Ann Metskas, Robert C. Hurt, Hossein Salahshoor, Ting-Yu Wang, Dina Malounda, George Lu, Tsui-Fen Chou, Mikhail G. Shapiro, Grant J. Jensen

## Abstract

Gas vesicles (GVs) are gas-filled protein nanostructures employed by several species of bacteria and archaea as flotation devices to enable access to optimal light and nutrients. The unique physical properties of GVs have led to their use as genetically-encodable contrast agents for ultrasound and MRI. Currently, however, the structure and assembly mechanism of GVs remain unknown. Here we employ cryo-electron tomography to reveal how the GV shell is formed by a helical filament of highly conserved GvpA subunits. This filament changes polarity at the center of the GV cylinder—a site that may act as an elongation center. High-resolution subtomogram averaging reveals a corrugated pattern of the shell arising from polymerization of GvpA into a β-sheet. The accessory protein GvpC forms a helical cage around the GvpA shell, providing structural reinforcement. Together, our results help explain the remarkable mechanical properties of GVs and their ability to adopt different diameters and shapes.

## INTRODUCTION

A fundamental property of many living organisms is their ability to move within their environment, with single-celled organisms capable of swimming, crawling and aligning with magnetic fields. The molecular machines underlying many of these motility functions have been characterized in detail (Komeili et al. 2006; Krause et al. 2018; Wadhwa and Berg 2022). Yet the structure underlying one of the oldest evolved forms of motility–flotation–remains more mysterious. Some cyanobacteria, heterotrophic bacteria, and archaea regulate their buoyancy in aquatic environments to access sunlight and nutrients using intracellular flotation devices called gas vesicles (GVs) (Walsby 1994; Pfeifer 2012). These unique protein nanostructures consist of a gas-filled compartment, typically ∼100 nm in diameter and ∼500 nm in length, enclosed by a ∼3 nm-thick protein shell (Figure 1A) that can withstand hundreds of kPa of applied pressure (Lakshmanan et al. 2017; Dutka et al. 2021). The interior of the shell is strongly hydrophobic, keeping out water while allowing gas molecules to diffuse in and out on a sub-millisecond timescale (Offner et al. 1998; Pfeifer 2012). By controlling the expression of GVs, cells alter their density, to move up or down in a column of water.

**Figure 1.**
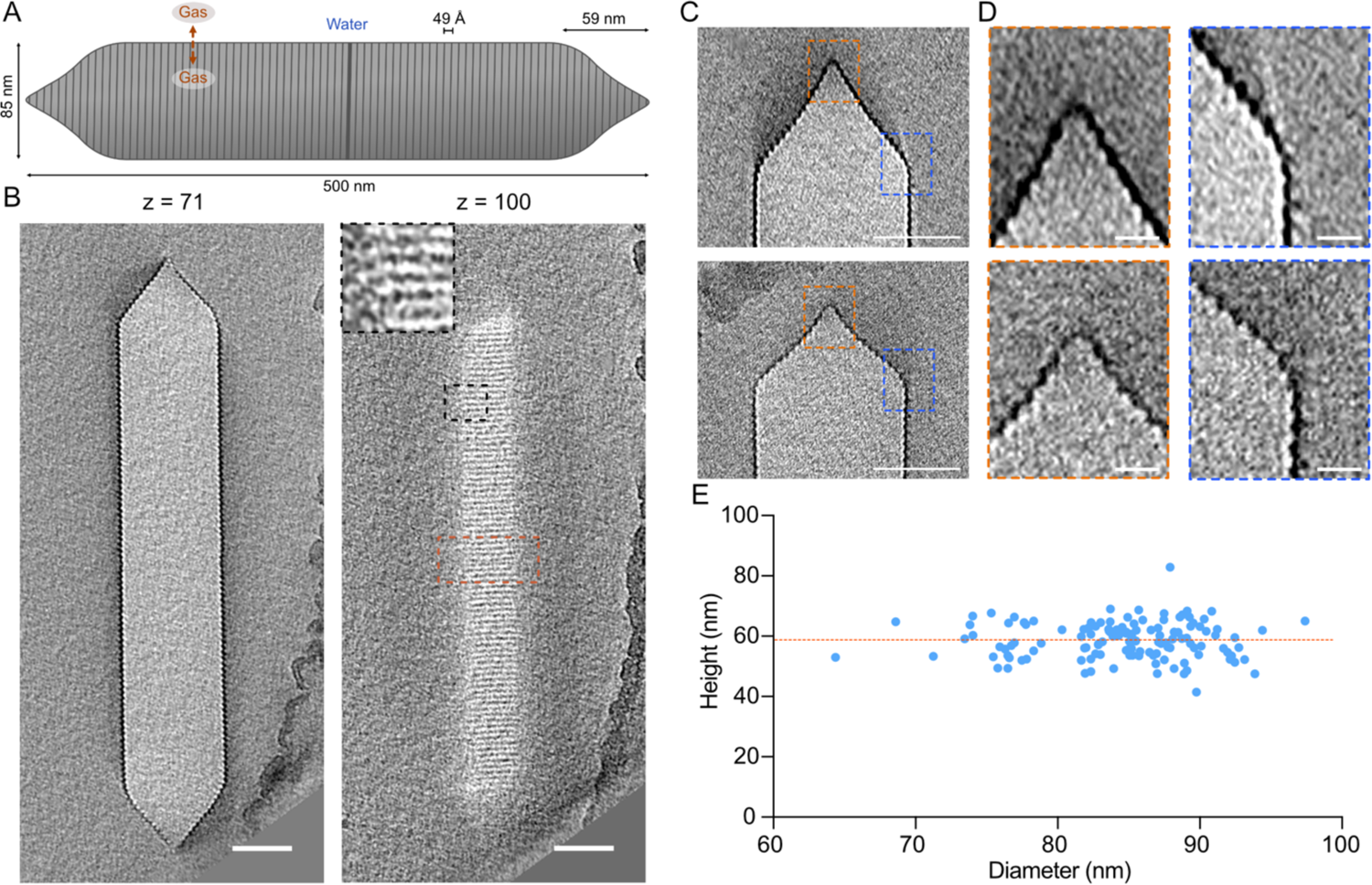
Molecular architecture of Ana GVs. (**A**) Schematic representation of an Ana GV with average dimensions annotated. (**B**) Representative slices at the indicated z-heights from a cryo-electron tomogram of an individual GV. Inset shows an enlargement of the area indicated by the black dashed box. Scale bars, 50 nm. (**C**) Central tomographic slices of two conical GV ends with different morphologies. Scale bars, 50 nm. (**D**) Enlarged views of the areas indicated by orange (apex) and blue (cone to cylinder transition) dashed boxes in **C**. Scale bars, 10 nm. (**E**) Distribution of the diameters and heights of conical GV ends, n= 132. Orange dashed line indicate average height of the cones (59 ± 6 nm).

In addition to their biological significance, GVs are a subject of intense interest for biotechnology. Analogously to fluorescent proteins, opsins and CRISPR nucleases, GVs’ unusual biophysical properties can be harnessed for other purposes. The gaseous composition of GVs allows them to scatter ultrasound waves, enabling their use as genetically-encoded reporters and actuators of cellular function deep in tissues (Shapiro, Goodwill, et al. 2014; Bourdeau et al. 2018; Farhadi et al. 2019; Wu et al. 2019; Farhadi et al. 2020; Lakshmanan et al. 2020; Bar-Zion et al. 2021; Hurt et al. 2021). Other applications take advantage of GVs’ refractive index, gas permeability and susceptibility to magnetic fields (Shapiro et al. 2014; Lu et al. 2018, 2020).

GVs were discovered in the 19^th^ century, but we still have limited knowledge of their structure and assembly. GVs adopt a cylindrical shape, with conical caps (Figure 1A). Their components are encoded in operons containing relatively few genes (8-23+, depending on the species) (Pfeifer 2012). One of these genes encodes the main structural protein, GvpA, a small (∼8 kDa), highly hydrophobic protein that polymerizes to form the GV shell (Walsby 1994). In some species, the gene cluster contains a secondary structural protein called GvpC, which binds to the exterior of the shell to provide mechanical reinforcement (Hayes et al. 1992). The remaining genes encode proteins whose functions are not well understood, possibly including chaperones, assembly factors, and additional minor shell constituents. GVs are nucleated as bicones which then elongate into a cylindrical shape with low-pitch helical ribs (Offner et al. 1998; Pfeifer 2012), but their detailed molecular structure is not known.

Here, we apply state-of-the-art cryo-electron tomography (cryo-ET) and sub-tomogram averaging techniques to GVs from the cyanobacterium *Anabaena flos-aquae* (Ana). These GVs are among the best studied by biophysicists (Walsby 1994; Maley et al. 2017; Cai et al. 2020) and the most commonly used in biotechnology applications (Lakshmanan et al. 2016, 2020; Hurt et al. 2021). We show that the Ana GV shell is formed by a continuous helical filament of repeating GvpA subunits, giving rise to a corrugated cylindrical structure with terminal cones that taper over a conserved distance. Near the middle of the cylinder, the angle of corrugation is inverted, suggesting a potential elongation center for GV biosynthesis. The corrugated shell is externally reinforced by circumferential rods of GvpC. Combining our cryo-ET data with an atomic model of the homologous *Bacillus megaterium* (Mega) GvpA protein determined in a complementary study (Huber et al. 2022), we build an atomic-level structural model of the Ana GV. This model explains the strong hydrophobicity of the GV interior and the connection between the GV shell and GvpC and highlights the structural conservation of GVs between diverse species. Finally, we extend our study with biochemistry and computational modeling to corroborate our model and explore its implications for GV engineering.

## RESULTS

### Molecular architecture of GVs

Ana GVs are long, cone-tipped cylinders with diameters of 85 ± 4 nm (Dutka et al. 2021) and lengths of 519 ± 160 nm (Lakshmanan et al. 2017) (Figure 1A and B). Although GVs have apparent helical symmetry, they are prone to deformations in thin ice (Figure S1) and are therefore intractable for cryo-EM helical processing. For this reason, we decided to use cryo-ET. However, cryo-ET analysis of GVs presents its own challenges. We observed that GVs are highly sensitive to electron dose, losing high-resolution features quickly before deflating and shrinking (Movie S1). To mitigate this effect, we limited the total electron dose to ∼45 electrons/Å^2^ per tilt-series, which is ∼2.5 times lower than typically used for high-resolution sub-tomogram averaging (Peukes et al. 2020; Metskas et al. 2022).

We started by examining large-scale structural features. While the diameter and length of GVs have been characterized (Walsby and Bleything 1988; Dutka et al. 2021), the conical ends and their connection to the cylindrical body are less studied. Close inspection of individual caps in our cryo-tomograms revealed a heterogenous morphology that deviated from simple conical structure (Figure 1C and D). We observed two elements in the majority of cones: a pointed closed tip, and a rounded transition region between cone and cylinder (Figure 1D). The height of the conical caps was 59 ± 6 nm, independent of cylinder diameter (Figure 1E). The rounding of the base was more pronounced in GVs with larger diameters, so we also examined cryo-tomograms of Mega GVs, whose average diameter is ∼30 nm smaller than that of Ana GVs. However, Mega GVs showed similar rounding at the cap transition (Figure S2), suggesting that this is a conserved feature of the structure independent of width.

### The GvpA spiral reverses polarity in the middle of the cylinder

The GV shell consists of a low-pitch helix, running the length of the GV (Figure 2A and 2B). Near the middle of the GV, however, the angle of the helix abruptly inverted. Previously, Waaland and Branton (Waaland and Branton 1969) noticed that one rib in the middle of the GV cylinder appears to be thicker than the others and suggested that this could be the growth point, where new GvpA subunits are added. Indeed, this abnormal rib was clearly visible in our tomograms (Figure 2A). To obtain a better understanding of the rib architecture in that region, we applied subtomogram averaging, which revealed that the angle of corrugation is opposite above and below the central rib (Figure 2B). This polarity inversion occurs within one rib, and the continuity of the spiral is not broken (Figure 2B and C). We were unable to distinguish whether the polarity of GvpA subunits changed relatively gradually within the space of one helical turn, or abruptly from one monomer to the next. We also could not tell whether additional proteins are present at the inversion point.

**Figure 2.**
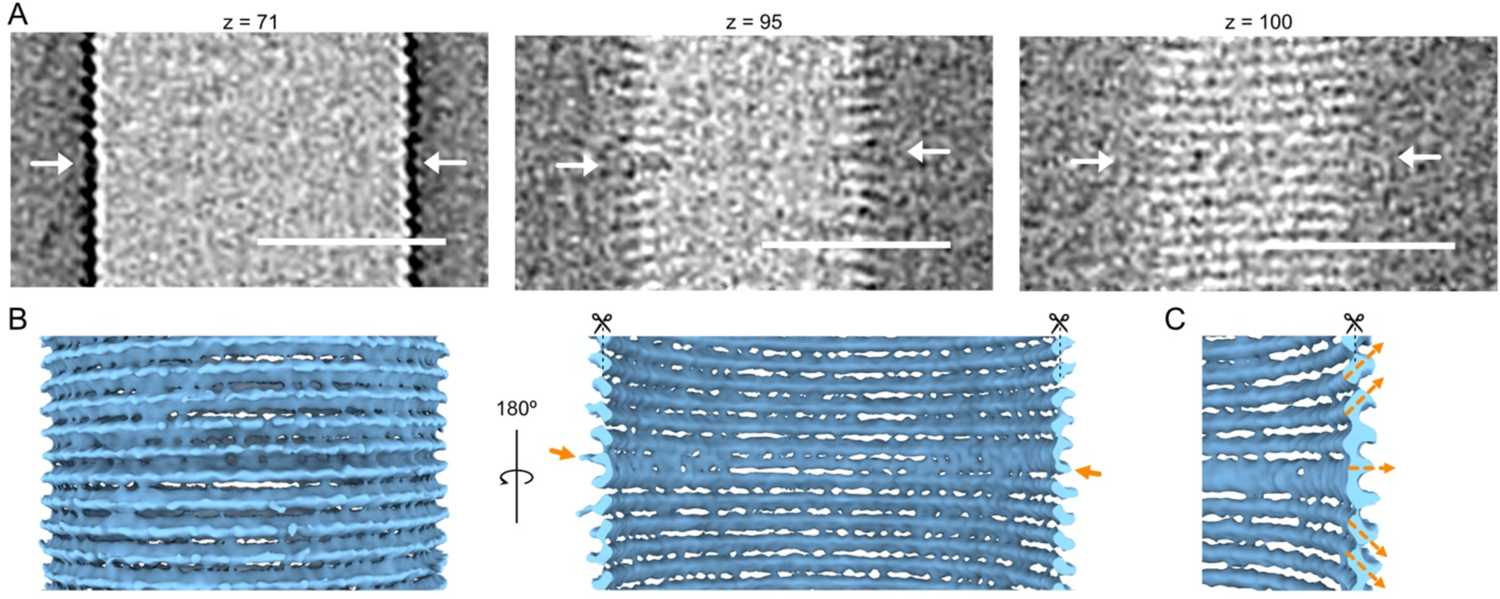
Polarity inversion point. (**A**) Tomographic slices at the indicated z-heights of the GV section indicated by the orange dashed box in Figure 1B. Scale bars, 50 nm. (**B**) Subtomogram average of the middle region of the GV where the ribs reverse polarity. Arrows denote the rib where polarity is reversed. (**C**) Enlarged view of the subtomogram average in (**B**) highlighting the inversion of the helical assembly.

By inspecting hundreds of cryo-electron micrographs of GVs from different species (*A. flosaquae, B. megaterium* and *Halobacterium salinarum*) we found that the polarity inversion point is a conserved feature (Figure S3). Although in general the inversion point was near the middle of the cylinder, in some cases it was located closer to one end (Figure S3A). If it is the nucleation point, this suggests that GvpA subunits are not always added symmetrically in both directions.

Additionally, we observed some examples where a GV exhibited different diameters on either side of the inversion point (Figure S3B). While we saw examples in all three species, it was most frequent, and most pronounced, in GVs from *H. salinarum* (Halo).

### Sub-tomogram averaging of the GV shell

To understand the molecular details of the GV structure, we applied subtomogram averaging to the Ana GV shell, both in its native state and after biochemically removing the reinforcing protein GvpC to produce “stripped” (AnaS) GVs. Initially, we tried averaging tubular sections of the GVs. However, due to flattening and the low number of particles, the resolution of this approach was limited (Figure 3A). As an alternative, we decided to average only small sections of the shell using an oversampling method (Peukes et al. 2020; Wan et al. 2020). This strategy produced a higher number of particles and allowed more rigorous 3D classification to remove distorted particles. With this method, we produced subtomogram averages of native Ana (Figure 3B) and AnaS (Figure 3C) GV shells with global resolutions of 7.7 Å and 7.3 Å, respectively (Table S1 and Figure S4). Both density maps showed a similar range of local resolution, with the best-resolved parts reaching 6.5 Å (Figure S4A and C). In both cases, the structure revealed a prominent pattern of beveled ribs, giving rise to the corrugated GV shell. The shell was ∼4 nm wide at its thickest and only ∼1 nm thick in the region between adjacent ribs (Figure 3D). We also observed pores in this region, at the interface between neighboring ribs of the spirals (Figure 3B and C), likely allowing gas to diffuse in and out of the GV. In contrast to the complex exterior face of the GV shell, the gas-facing interior appeared relatively smooth.

**Figure 3.**
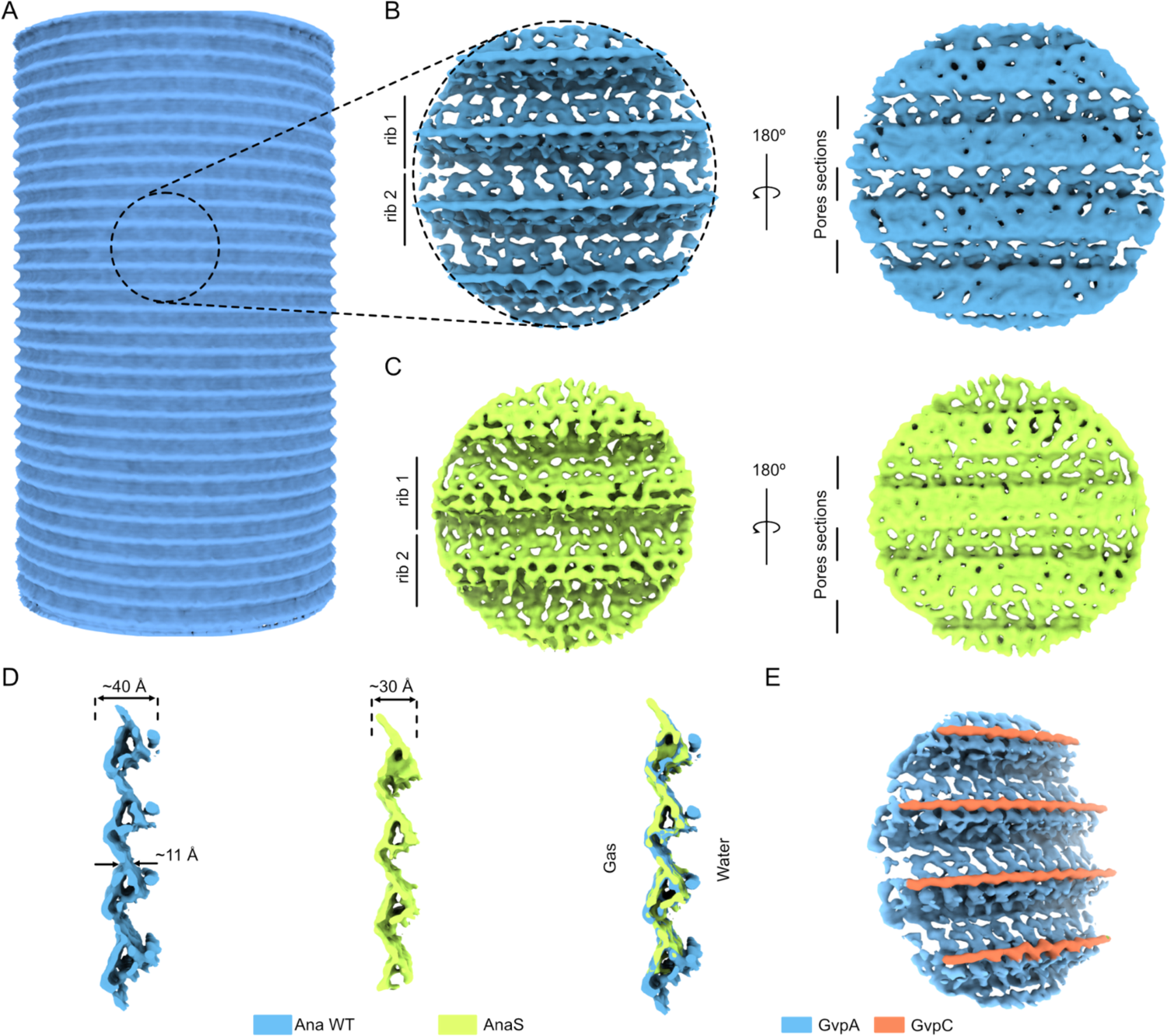
Cryo-ET structure of the Ana GV shell. (**A**) Initial, low-resolution subtomogram average of a cylindrical GV segment. (**B**) Orthogonal views of a higher-resolution (7.7 Å) sub-tomogram average of the native Ana GV shell. (**C**) Orthogonal views of a higher-resolution (7.3 Å) sub-tomogram average of the AnaS GV shell. (**D**) Cross-sections of the subtomogram averages of the GV shell, superimposed at right. (**E**) Segmented density map of the native Ana GV indicating the locations of GvpC.

Comparing the maps of native Ana and AnaS GVs (lacking GvpC), we noticed a pronounced rod-like structure positioned along the GV ribs that is absent in AnaS (Figure 3D). Previously, various models for GvpC binding to the GV shell have been proposed (Buchholz, Hayes, and Walsby 1993), with most of the field favoring one in which GvpC spans longitudinally across GvpA ribs (Maresca et al. 2018; Lakshmanan et al. 2020). Our structure shows instead that GvpC binds circumferentially to the thickest part of the GV shell, thereby creating a spiral cage around the GV cylinder (Figure 3E). We do not yet know whether the GvpC filament binds the central inversion rib or extends to the conical caps, where the decreasing radius of curvature might be prohibitive, or whether it is continuous, as the average would blur away gaps.

### GvpA polymerizes into a β-sheet to form GV ribs

The resolution of our Ana GV density map was sufficient for rigid-body fitting of a homology model of GvpA. Taking advantage of the high degree of conservation of the protein, we used the structure of GvpA1 from *B. megaterium* solved by helical reconstruction in a contemporaneous study (Huber et al. 2022). The only substantial difference between GvpA from Ana and Mega is an extended C-terminus in the latter (Figure S5), so our homology model was complete and fit well into our cryo-ET density map (Figure 4A). In agreement with a previous NMR prediction, we saw that individual GvpA subunits adopt a coil-α-β-β-α-coil motif (Sivertsen et al. 2010) (Figure 4B). Adjacent subunits form antiparallel β-sheets, and N-terminal coils mediate interactions between ribs, extending toward the β-sheet of subunits above and then out towards their C-terminal α2 helices. All domains of the small GvpA protein play a role in building the GV shell, packing into a tight structure with only small pores. The outer surface of the GV was largely hydrophilic, with small hydrophobic pockets between the α2 helices, but the interior was strongly hydrophobic (Figure 4C), explaining how GVs can prevent the entry or condensation of liquid water, a property that differentiates them from all other known protein assemblies. This combination of extreme amphiphilicity and tight interactions between GvpA subunits also explains the remarkable stability of GVs; we find purified GVs are stable for years at cool or ambient temperature.

**Figure 4.**
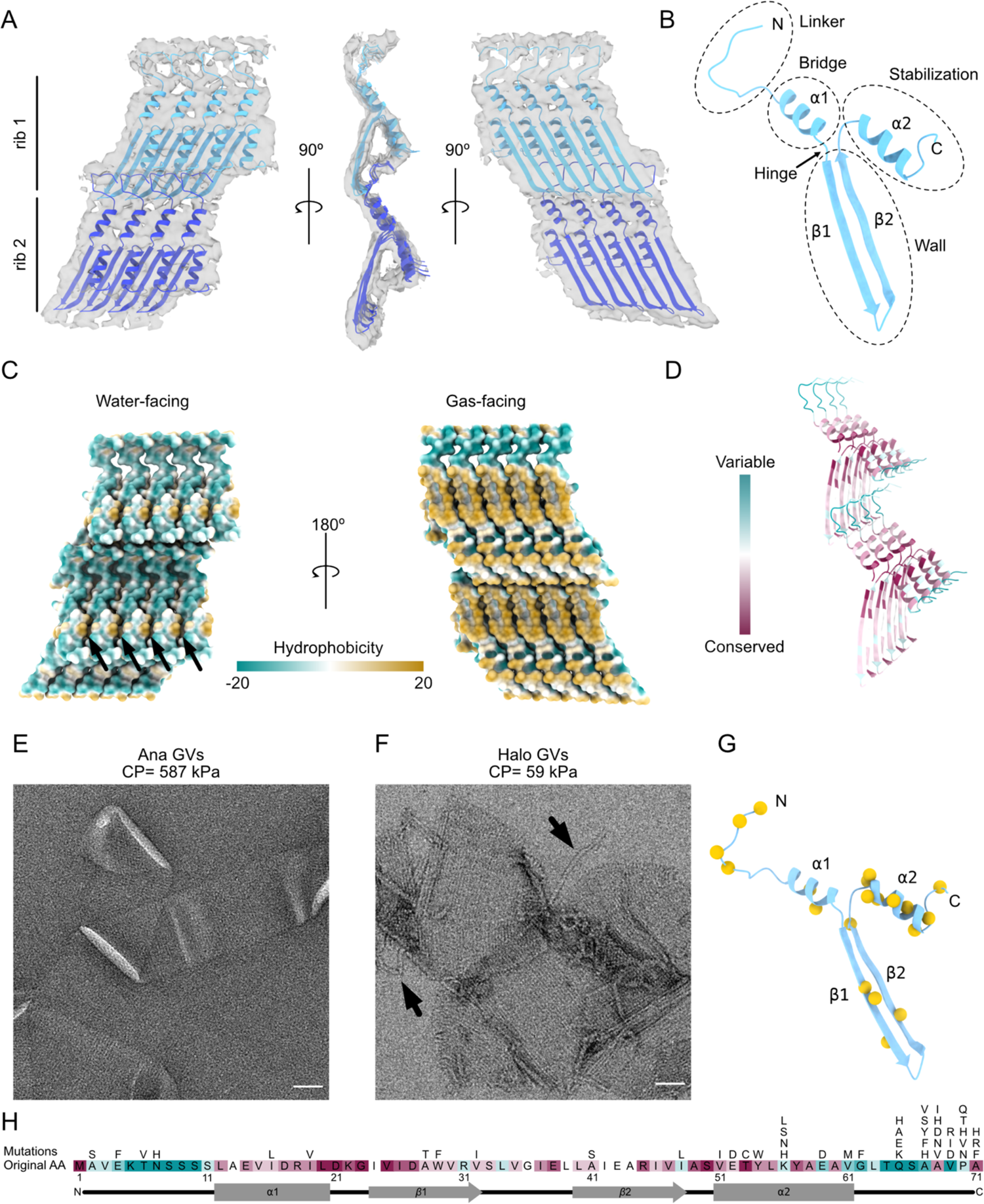
Atomic model of the Ana GV shell. (**A**) 7.3 Å resolution structure of two adjacent GvpA ribs determined by subtomogram averaging (grey surface), fitted with a homology model of GvpA. (**B**) An individual GvpA subunit with domains annotated. (**C**) Surface views of the exterior and interior faces of the GV shell, colored by hydrophobicity. Arrows indicate hydrophobic pockets located between α2 helices. (**D**) Conservation analysis of GvpA determined by ConSurf (Ashkenazy et al. 2016). (**E, F**) Negative-stain EM images of collapsed GVs from (**E**) *A.flos-aquae* and (**F**) *H.salinarum*. Arrows indicate separated GvpA filaments. Collapse Pressure (CP) is indicated above. Scale bars, 50 nm. (**G**) Location of tolerated mutation sites (yellow spheres) in the GvpA structure (blue). (**H**) Map of all tolerated mutations in GvpA. Original sequence colored by conservation score as in **D**.

As mentioned above, the only major difference between *B. megaterium* GvpA1 and *A. flos-aquae* GvpA is the presence of an elongated C-terminus (Figure S5). This C-terminus was not resolved in a recent structure solved by helical processing (Huber et al. 2022), presumably due to its flexibility. In our cryo-tomograms of Mega GVs, we observed additional density on the surface of the shell (Figure S6A-C) that is absent from the structures of AnaS and native Ana GV shells (Figure S6D and E). The density was not highly regular but appeared connected. It may be that this extra density belongs to the C-terminus of GvpA1, which perhaps plays a role in stabilizing the GV shell.

The sequence of GvpA, the major structural protein, is highly conserved in all GV-producing species (Englert, Horne, and Pfeifer 1990; Griffiths, Walsby, and Hayes 1992) and we think it likely that its structure is similarly conserved, as evidenced by a model from *B. megaterium* GvpA1 (Huber et al. 2022) and fitting into the density of *A. flos-aquae* GvpA. Remarkably, though, GvpA can assemble into GVs with varying diameters (Figure S7A) (Dutka et al. 2021) and morphologies (Figure S7B and S7C). For instance, the largest Halo GVs are ∼7-times larger in diameter than the smallest Mega GVs. One key to understanding different morphologies may lie in what appears to be a hinge region located between helix α1 and strand β1 (Figure 4B), where a conserved glycine resides (Figure 4H and S5). Small sequence differences in GvpA have been suggested to contribute to different morphologies of GVs (Walsby 1994). *H. salinarum* contains two independent GV gene clusters, p-vac and c-vac (Pfeifer 2012). The sequences of the GvpA homologs in the two clusters are 94% identical (Figure S5), yet GVs produced by the c-vac gene cluster adopt a lemon shape (Figure S7B) while p-vac produces the more typical cylindrical shape with conical caps (Figure S7C).

We used ConSurf (Ashkenazy et al. 2016) to visualize the evolutionary conservation of GvpA, revealing that the most conserved residues are located in the β-sheets and α-helices (Figure 4D). In contrast, the N-terminal domains of the protein responsible for interactions between neighboring ribs showed the greatest variability (Figure 4D). Within the generally conserved β-strands, the most variable sites were those interacting with the N-terminus from the subunit below. This variability in amino acid composition in the domains responsible for holding adjacent ribs together might be one factor contributing to differences in the mechanical strength of GVs. Under hydrostatic pressure, GVs can collapse, forming flattened sacs (Dutka et al. 2021). The critical pressure required to collapse GVs varies greatly between species. For example, the hydrostatic collapse pressure threshold of Ana GVs is 587 kPa, while that of Halo GVs is 59 kPa, an order of magnitude lower (Lakshmanan et al. 2017). By EM imaging, we found that Ana GVs collapse without major disruptions to the rib structure (Figure 4E), while collapsed Halo GVs often exhibit major disruption of the rib structure and separation of the GvpA filament (Figure 4F). This supports the idea that the strength of connectivity between ribs varies between species.

To test the importance of conserved GvpA residues in GV assembly, we mapped tolerated mutations by screening a scanning site-saturation library of GvpA mutants in *Escherichia coli* engineered to express a hybrid gene cluster composed of the structural proteins GvpA and GvpC from the Ana GV gene cluster and the assembly factors GvpR-GvpU from the Mega GV gene cluster. GV-producing mutant clones were identified by nonlinear x-wave ultrasound (xAM) (Figures 4G,H, and S8). The results largely correlated with observed evolutionary conservation, with the highest number of function-retaining mutations occurring in the evolutionarily variable C-terminal coil (Figure 4H). Interestingly, the only conserved region that tolerated mutations well was helix α2, which is not involved in interactions between monomers, but rather plays a crucial role in GvpC binding (see below).

### GvpC forms a helical spiral around the GV shell

Having identified GvpC in our subtomogram average of the Ana GV shell (Figure 3E), we next investigated how GvpC binds to GvpA and how multiple GvpC proteins might cooperate to strengthen GVs. GvpC is predicted to form an amphipathic α-helical structure composed of a characteristic 33-residue repeating sequence (Figure S9A). *A. flos-aquae* GvpC consists of 5 such repeats, plus short N- and C-termini. To build a model of GvpA decorated with GvpC, we fitted the AlphaFold2 (Mirdita et al. 2022) structure prediction for the third repeat of *A. flos-aquae* GvpC into our subtomogram average. We first fitted the predicted model in an orientation where the hydrophobic site of the helix was facing the GV shell, and then subsequently refined the placement to optimize geometry and reduce clashes (Figure 5A).

**Figure 5.**
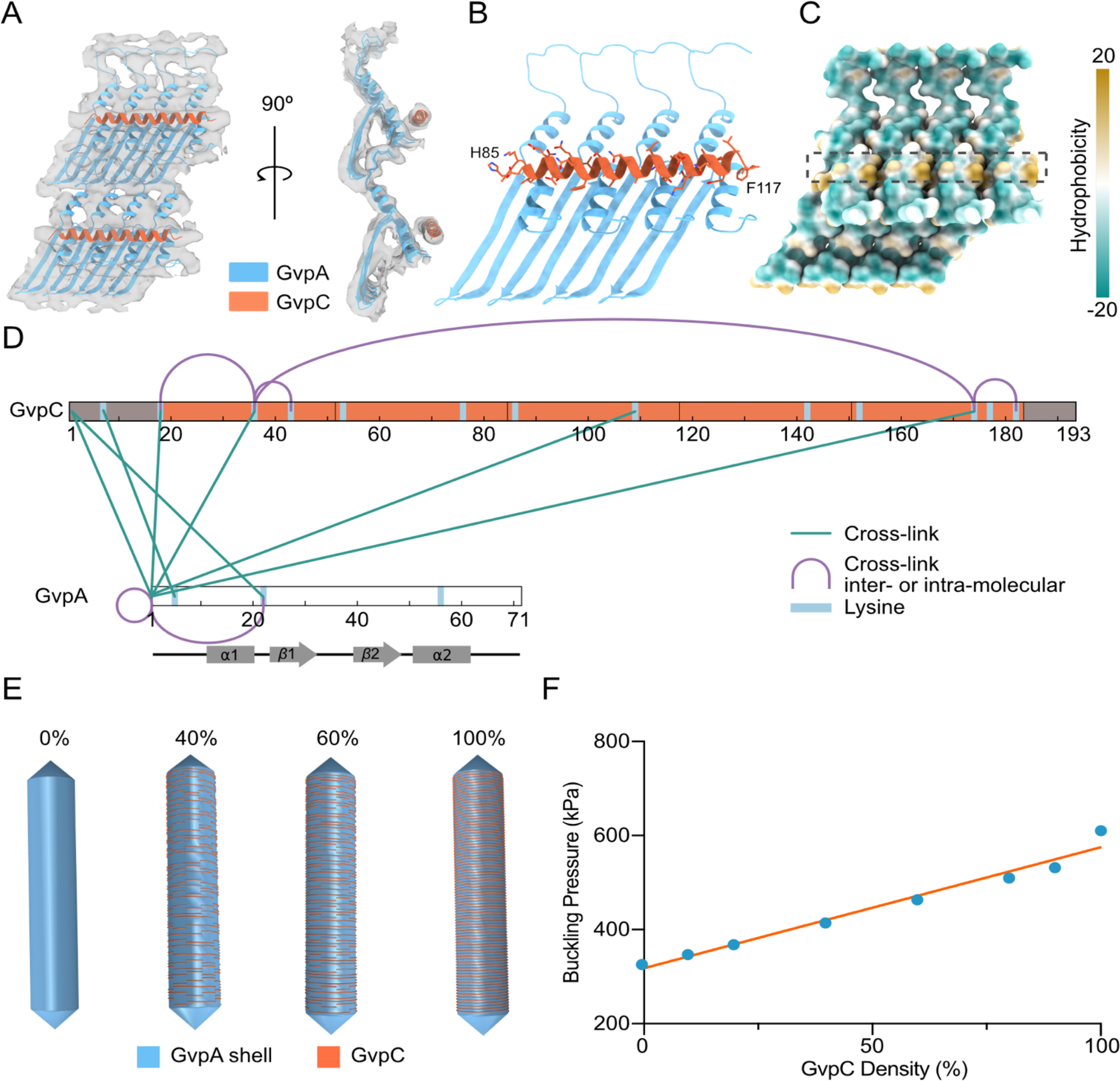
Mechanical reinforcement of the GV shell. (**A**) 7.6 Å resolution subtomogram average of neighboring Ana GvpA monomers connected by GvpC (grey surface) fitted with a homology model of GvpA and an AlphaFold2 model of the third repeat of GvpC. (**B**) Resulting GvpC binding model. (**C**) GvpC binding site (dashed black box) at the hydrophobic pockets between α2 helices of GvpA. The surface of GvpA is colored by hydrophobicity. (**D**) Crosslinked sites between GvpA and GvpC identified by mass spectrometry. (**E**) Finite element shell models of a GV with a length of 500 nm and width of 85 nm and the indicated degree of GvpC saturation. (**F**) Buckling pressure as a function of GvpC density. The orange line represents a simple linear regression fit.

We found that GvpC binds perpendicularly to the surface-exposed α2 helices of GvpA, directly above the hydrophobic pockets (Figure 5A-5C and 4C). The bulky and hydrophobic residues are located mainly between helices α2 of GvpA (Figure 5B). In addition to being amphipathic, GvpC also has an unequal distribution of charge (Figure S9B). In our model, GvpC binds directly above the negatively-charged C-terminus of GvpA (Figure S9C). One molecule of GvpC (not including the N- and C-termini) interacts with approximately four GvpAs, indicating a GvpC to GvpA ratio of at least 1:20 if saturated. This is close to the previously calculated ratio of 1:25 (Buchholz, Hayes, and Walsby 1993). In our model, we were not able to confidently predict the directionality of GvpC relative to the GvpA spiral, but attempts to refine GvpC in the reverse orientation in Phenix resulted in worse refinement scores.

Despite multiple rounds of 3D classification and application of different focus masks, we were unable to resolve the junctions between neighboring GvpC molecules. Instead, GvpC appeared as a continuous helical belt. To get a better understanding of GvpC-GvpC and GvpC-GvpA interactions, we performed chemical cross-linking coupled with mass spectrometry (XLMS) (Table S2). Most of the cross-links we observed were between the N-terminus of GvpA and apparently random locations on GvpC (Figure 5D), which is consistent with the close association between the N-terminus of GvpA and the GvpA α2 helix in the adjacent rib, where GvpC binds, in our structure (Figure 5A). However, we did not observe any cross-links between GvpC and helix α2, potentially due to the unfavorable orientation of the lysines. Among GvpC-GvpC cross-links, the most interesting was between K36 and K174 (Figure 5D). The distance between these residues is ∼20 nm, too far for an intramolecular cross-link (Merkley et al. 2014), suggesting that GvpC termini are either closely packed or potentially interact tail-to-tail (Figure S10).

To quantify the effect of increasing GvpC occupancy on GV stabilization, we used solid mechanics simulations to estimate the applied pressure at which the GV shell starts to buckle–a parameter relevant to its ability to withstand hydrostatic pressure, as well as produce nonlinear signal in ultrasound imaging. We implemented several finite element models of a GV shell, each 500 nm in length and 85 nm in diameter, and with a custom density of GvpC molecules. From a continuous belt, representing 100% GvpC, we randomly removed GvpC-length (25 nm) segments of the helix to achieve the desired saturation for each model (Figure 5E). We subjected the outer surface of each GV shell to uniform normal stress, simulating hydrostatic or acoustic pressure, and obtained a critical buckling pressure by linear buckling analysis. We observed a simple linear dependence of buckling on scaffolding protein density (Figure 5F), consistent with previous experimental findings that GvpC level can be utilized to modulate the GV buckling threshold (Lakshmanan et al. 2016)

## DISCUSSION

The GV shell has remarkable mechanical properties: despite being only ∼3 nm thick, it is highly stable and can withstand up to hundreds of kPa of pressure. This is achieved by tight packing of the GvpA subunits into a low pitch helix that forms a corrugated cylinder. On the macroscopic level, corrugation is typically used when flexibility is important (*e.g*., pipes) or to increase durability and strength (*e.g*., unpressurized cans). One or both of these properties might be similarly important for GV function. Our data indicate that GV cylinders can be significantly deformed without collapsing the structure (Dutka et al. 2021). This elasticity of the GV shell may be crucial for adapting to pressure fluctuations *in vivo*, and enables GVs to be used as contrast agents in high-specificity nonlinear ultrasound imaging (Maresca et al. 2017). We noticed a highly-conserved glycine between helix α1 and strand β1 of GvpA. The single hydrogen in the side chain of glycine gives it much more flexibility than other amino acids (Huang and Huang 2018), suggesting that this region may act as a hinge that confers elasticity on the shell structure and lets it adapt to different geometries, such as those observed in terminal cones or the bodies of lemon-shaped GVs.

The primary contact between adjacent GvpA subunits is mediated by lateral interactions of antiparallel β-strands in an extended sheet, resembling the aggregation of β-amyloids (Liberta et al. 2019; Berhanu et al. 2015). Such assemblies are typically stabilized by an extensive network of backbone hydrogen bonding, conferring outstanding strength (Paul et al. 2016). Such strength is also observed in GVs from diverse species; individual GvpA monomers can only be dissociated from the polymer by harsh chemical treatment (Walker and Walsby 1983; Belenky et al. 2004). That backbone interactions are the main force driving subunit polymerization is consistent with the wide range of diameters observed in different species (Dutka et al. 2021): as the curvature of the cylinder changes, the relative positions of backbone residues will be affected much less than those of side chains. We find that GvpA domains involved in forming the GV wall have a low tolerance for mutations, likely due to selective pressure to preserve the highly hydrophobic composition of the β-sheets and maintain interactions with the linker domain connecting subsequent coils of the helix.

Stacked ribs of the continuous GvpA polymer are joined by interactions of the coiled N-termini from one row of subunits with the β-strands of the subunits in the next. We observe that the strength of these inter-rib interactions varies between species, likely related to evolutionary variability in the N-terminal linker. It was previously observed that the critical collapse pressure of Mega GVs is much higher than that of Ana or Halo GVs (Lakshmanan et al. 2017), likely due to the narrower diameter of Mega GVs (Beard et al. 1999, 2000). However, we note that the C-terminus of Mega GvpA is longer than in other species and in our tomograms of Mega GVs, we observed extended irregular surface densities connecting ribs. We suggest that these extra densities correspond to the extended C-termini of *B.megaterium* GvpA1 and may confer additional mechanical strength.

Other mechanisms also enhance the strength of the GV shell. Almost all GV gene clusters encode an additional, minor structural protein, GvpC, that binds to the GvpA helical spiral and reinforces the shell (Walsby and Hayes 1988; Lakshmanan et al. 2016); we find that GvpC binds to the surface-exposed α2 helix of GvpA. In our mutational analysis, this helix was relatively mutation-tolerant, suggesting that it has a minimal role in GvpA shell integrity and instead acts primarily as an adapter for GvpC. In contrast to previous models of GvpC spanning ribs, we find that GvpC instead tracks along ribs, forming a spiral cage around the GV cylinder. Our XLMS results indicate close conjunction of GvpC molecules and, even with multiple masking and 3D classification strategies, we never observed discontinuity in the GvpC rod in our subtomogram averages. Although we could not resolve interactions between GvpC N- and C-termini, we previously showed that their removal leads to a significant drop in critical collapse pressure of Ana GVs (Lakshmanan et al. 2016). Here, we used finite element simulations to quantify the reinforcing effect of GvpC density on GV buckling and find that the degree of strengthening is directly proportional to the amount of GvpC bound. However, full GvpC occupancy is not required for full strengthening, and small gaps in the GvpC cage have a negligible effect on collapse pressure.

In the initial stage of assembly, GVs grow as bicones until reaching their target diameter; at that point, growth elongates the central section, producing cylinders which can reach several micrometers in length (Pfeifer 2012; Farhadi et al. 2019). The trigger for this transition is unclear. Our data show that the height of mature cones is relatively constant, regardless of GV diameter, indicating that the number of helical turns/height is the measured quantity, rather than the number of GvpA subunits. Our observation of a polarity inversion near the middle of the GV suggests that this is the site of cylinder elongation, with individual subunits being incorporated in both directions. In some cases, we observed that the elongation center was located closer to one end of the GV, suggesting a mechanism that does not require GvpA subunits to be added symmetrically in both directions. Although GV cylinders typically exhibit a uniform diameter, we documented some examples with different diameters on either side of the elongation center. We observed variations in the shape of conical ends, both within and between GVs. This hints that mismatches in GV geometry might arise in the initial bicone growth stage, but further investigation is needed to fully dissect the mechanism of GV morphogenesis.

Currently, the method of choice for solving the structure of helical assemblies is helical reconstruction (Egelman 2015; He and Scheres 2017). However, the large and nonuniform diameter of GVs and their susceptibility to deformation during cryopreservation present challenges for this approach. Cryo-ET and subtomogram averaging can circumvent these limitations by focusing on smaller, and therefore more uniform, 3D sections of the object of interest. Subtomogram averaging can reach high resolution in certain favorable cases such as for large (Tegunov et al. 2021) or symmetrical (Schur et al. 2016; Metskas et al. 2022) proteins, but for most targets, resolution has remained limited. Here we show that even with a fairly challenging target, recent developments in cryo-ET data collection and subtomogram averaging methods make it possible to obtain sufficient resolution to unambiguously dock an atomic model. Our work, together with a complementary study of Mega GVs (Huber et al. 2022), advances our understanding of the molecular architecture of GVs and may inform further engineering of GVs to serve as genetically-encoded contrast agents and biosensors.

## MATERIAL AND METHODS

### GV preparation

GVs were isolated either from native sources, *Anabaena flos-aquae* (Ana) and *Halobacterium salinarum* NRC1 (Halo), or expressed heterologously in Rosetta 2(DE3)pLysS *Escherichia coli, Bacillus megaterium* (Mega), as previously described (Lakshmanan et al. 2017). In the final steps of buoyancy purification, the sample buffer was exchanged for 10 mM HEPES, pH 7.5. To obtain GVs stripped of GvpC (AnaS), 6 M urea solution was added to purified native GVs and two additional rounds of buoyancy purification were performed. AnaS GVs were subsequently dialyzed in 10 mM HEPES, pH 7.5. Concentrations were measured by optical density (OD) at 500 nm using a spectrophotometer (NanoDrop ND-1000, Thermo Scientific).

### Cryo-ET

A freshly purified GV sample was diluted to OD_500_ = ∼20 (Ana and Halo), ∼3 (AnaS), or ∼1 (Mega) and mixed with 10 nm BSA-coated gold beads. A 3 μL volume of sample was applied to C-Flat 2/2 - 3C grids (Protochips) that were freshly glow-discharged (Pelco EasiGlow, 10 mA, 1 min). GV samples were frozen using a Mark IV Vitrobot (FEI, now Thermo Fisher Scientific) (4°C, 100% humidity, blot force 3, blot time 4 s).

Tilt-series were collected on a 300 kV Titan Krios microscope (Thermo Fisher Scientific) equipped with a K3 6k × 4k direct electron detector (Gatan). Multi-frame images were collected using SerialEM 3.39 software (Mastronarde 2005) using a dose-symmetric tilt scheme. Super-resolution movies were acquired at a pixel size of 0.8435 Å (53,000× magnification) with varying defocus from − 1.0 to − 3.5 μm. Tilt-series of Halo and Mega GVs were collected from −60° to 60° with 3° increments. Tilt-series of native Ana GVs were collected in two sessions. The first set was collected from −60° to 60° with 3° increments and the second from −44° to 44° with 4° increments. For AnaS GVs, data were collected from − 45° to 45° with 3° increments. Due to the rapid shrinking of GVs during exposure to the electron beam (Movie S1), the total accumulated dose in all cases was limited to 45 electrons/Å^2^. Data collection parameters are summarized in Table S1.

Raw movies were binned by a factor of 2 and gain- and motion-corrected on-the-fly using Warp (Tegunov and Cramer 2019). Assembled tilt-series were exported to Dynamo (Castaño-Díez et al. 2012) for automated alignment using *autoalign_dynamo* (Burt et al. 2021). Aligned tilt-series were CTF corrected and full tomograms were either reconstructed in Warp with a pixel size of 10 Å or manually aligned and reconstructed using Etomo (Mastronarde and Held 2017).

### Subtomogram averaging - inversion point

Sub-volume extraction, alignment, and averaging were performed using the Dynamo software package (Castaño-Díez et al. 2012). Particles for subtomogram averaging of the inversion site were manually selected from GVs with a diameter of ∼85 nm, yielding a total of 68 particles. Sub-volumes were extracted from 4x binned tomograms with a final pixel size of 6.748 Å and 180×180×180 box size. The initial reference for particle alignment was generated by averaging segments with azimuth-randomized orientations. Due to the low number of particles, subtomogram averaging was not performed according to a gold standard. Instead, convergence of the structure was analyzed by changes in particle shifts and cross-correlation scores. During the final rounds of refinement, a soft cylindrical mask was applied to the central 40% of the GV tube.

### Subtomogram averaging - GV shell

Subtomogram averaging was carried out using Dynamo (Castaño-Díez et al. 2012), Warp (Tegunov and Cramer 2019), Relion-3.1 (Zivanov et al. 2018), and M (Tegunov et al. 2021) software packages. Data transfer between Dynamo and Warp/M was carried out using a set of tools provided by *warp2catalogue* and *dynamo2m* (Burt et al. 2021). Particle selection and initial reference generation were performed using the Dynamo package. Orientations and positions of shell sections were determined using geometrical tools for particle picking in Dynamo (Castaño-Díez, Kudryashev, and Stahlberg 2017). Initial estimates of positions and orientations on the GV shell were generated with an interparticle distance of ∼150 Å (∼3 ribs). Particles were extracted in Dynamo with a pixel size of 10 Å and averaged. After removal of duplicated particles, data was transferred to Warp and subtomograms were reconstructed with a pixel size of 5 Å based on the alignment information from Dynamo. Subtomograms were subsequently refined in RELION, re-reconstructed at 2.5 Å/pixel and 3D classified without alignment in RELION. After 3D classification, several additional rounds of 3D refinement were carried out in RELION. Finally, subtomograms were reconstructed at 1.687 Å/pixel and iteratively refined in RELION and M using a soft-edged mask around ∼3 or 4 adjacent ribs. Although we did not see a resolution boost after iterative refinement of the tilt-series parameters in M, subsequent refinement in RELION produced a better-quality reconstruction when applied to particles reconstructed after M refinement. Final maps were post-processed in RELION and resolution was estimated using a soft-edged mask around ∼3-4 adjacent ribs. The final results are summarized in Table S1.

### Model building

Although the density map for AnaS reached a higher overall resolution, individual features were better resolved in the map of native Ana GVs (Figure S4), so all model building was performed using this map. To build the GvpA model, a high-resolution cryo-EM structure of the homologous GvpA1 from *B. megaterium* (PDB:7R1C) (Huber et al. 2022) was fitted into the segmented cryo-ET density map corresponding to an individual subunit in UCSF Chimera (Pettersen et al. 2004). The GvpA amino acid sequence was rebuilt by manual replacement of mismatched residues in Coot (Emsley et al. 2010). The *A. flos-aquae* GvpA model was subsequently refined by rigid-body fitting using the Phenix real-space refinement tool (Adams et al. 2010). The refined GvpA model was used to populate a larger section of the cryo-ET map in UCSF Chimera (Pettersen et al. 2004). The Multimeric GvpA model was further refined by rigid-body fitting and refinement in Phenix to maximize fit into the density map. The model was inspected and manually optimized in Coot (Emsley et al. 2010) between subsequent refinement rounds in Phenix. The GvpC model was built by placing the AlphaFold2 (Jumper et al. 2021; Mirdita et al. 2022) structure prediction for the third repeat in an orientation where the hydrophobic side was facing the GV shell. The model was further refined to optimize helix geometry and minimize clashes in Phenix. The quality of the fit was evaluated using MolProbity (Chen et al. 2010).

### Cross-linking mass spectrometry (XLMS)

The cross-linking procedure was carried out according to the manufacturer’s instructions (Thermo Fisher). In brief, a freshly purified sample of native Ana GVs in 10 mM HEPES, pH 7.5 was mixed with an excess of cross-linker: either DSSO or BS3 (Thermo Fisher). The sample was incubated for 1h at room temperature and subsequently the reaction was quenched with Tris buffer at a final concentration of 20 mM.

The crosslinking samples were digested in an S-Trap mini spin column (Protifi, USA) according to the manufacturer’s instructions. For trypsin digestion, an additional aliquot of trypsin was added after 24 hours on the S-trap column and the digestion continued for another 24 hours. After elution and drying, peptides were suspended in LCMS-grade water containing 0.2% formic acid and 2% acetonitrile for further LC-MS/MS analysis. LC-MS/MS analysis was performed with an EASY-nLC 1200 (Thermo Fisher) coupled to a Q Exactive HF hybrid quadrupole-Orbitrap mass spectrometer (Thermo Fisher). Peptides were separated on an Aurora UHPLC Column (25 cm × 75 μm, 1.6 μm C18, AUR2-25075C18A, Ion Opticks) with a flow rate of 0.35 μL/min for a total duration of 43 min and ionized at 1.7 kV in the positive ion mode. The gradient was composed of 6% solvent B (2 min), 6-25% B (20.5 min), 25-40% B (7.5 min), and 40–98% B (13 min); solvent A: 2% ACN and 0.2% formic acid in water; solvent B: 80% ACN and 0.2% formic acid. MS1 scans were acquired at a resolution of 60,000 from 375 to 1500 m/z, AGC target 3e6, and a maximum injection time of 15 ms. The 12 most abundant ions in MS2 scans were acquired at a resolution of 30,000, AGC target 1e5, maximum injection time 60 ms, and normalized collision energy of 28. Dynamic exclusion was set to 30 s and ions with charges +1, +7, +8, and >+8 were excluded. The temperature of the ion transfer tube was 275°C and the S-lens RF level was set to 60. For cross-link identification, MS2 fragmentation spectra were searched and analyzed using Sequest and XlinkX node bundled into Proteome Discoverer (version 2.5, Thermo Scientific) against *in silico* tryptic digested *Dolichospermum-flos-aquae* GvpA from the Uniprot database. The maximum missed cleavages were set to 2. The maximum parental mass error was set to 10 ppm, and the MS2 mass tolerance was set to 0.05 Da. Variable crosslink modifications were set DSS (K and protein N-terminus, +138.068 Da) for BS3 crosslink and DSSO (K and protein N-terminus, +158.004 Da) for DSSO crosslink, respectively. For BS3 crosslink, the dynamic modifications were set to DSS hydrolyzed on lysine (K, +156.079 Da), oxidation on methionine (M, +15.995 Da), protein N-terminal Met-loss (−131.040 Da), and protein N-terminal acetylation (+42.011 Da). For the DSSO crosslink, the dynamic modifications were set to DSSO hydrolyzed on lysine (K, +176.014 Da), DSSO Tris on lysine (K, +279.078 Da), oxidation on methionine (M, +15.995 Da), protein N-terminal Met-loss (−131.040 Da) and protein N-terminal acetylation (+42.011 Da). Carbamidomethylation on cysteine (C, +57.021 Da) was set as a fixed modification. The false discovery rate (FDR) for crosslinked peptide validation was set to 0.01 using the XlinkX/PD Validator Node and crosslinks with an Xlinkx score greater than 30 were reported here. The raw data have been deposited to the ProteomeXchange Consortium (Deutsch, Bandeira, and Sharma, n.d.) via the PRIDE (Perez-Riverol et al. 2019) partner repository.

### Scanning site saturation library generation and screening

The scanning site saturation library was constructed via a Gibson assembly-based version of cassette mutagenesis as previously described (Ravikumar et al. 2018). Briefly, the *A. flos-aquae* GvpA coding sequence was divided into sections that tiled the gene, and oligos were designed to have a variable middle region with flanking constant regions against which PCR primers with Gibson overhangs were designed. The variable region was designed to sequentially saturate each residue with every amino acid other than the WT at that position, plus a stop codon to produce truncation mutants (*i.e*., the size of such libraries is 20 * [# of amino acids in the protein]). Oligos were synthesized as a pool by Twist Biosciences, and were amplified by 10 cycles of PCR (both to make them double-stranded and to add overhangs for Gibson assembly) using Q5 polymerase (according to the manufacturer’s protocol, but with 5 μM of each primer) and assembled with the rest of the GV gene cluster (*i.e*., Ana GvpC and Mega GvpR-GvpU) into a pET28a vector via Gibson assembly using reagents from New England Biolabs. Assembled libraries were electroporated into NEB Stable *E. coli* and grown in Lennox LB with 100 ug/mL kanamycin and 1% glucose (Ammar, Wang, and Rao 2018). Plasmid DNA was miniprepped (Econospin 96-well filter plate, Epoch Life Science) and verified by Sanger sequencing. Ultrasound-based phenotyping of mutants was performed in BL21-AI (Thermo) as previously described (Hurt et al., n.d.), and all screened mutants were sequenced using the evSeq pipeline (Wittmann et al. 2022).

### Finite element simulation

We first developed a finite element model of a single stripped GV isolated from *A. flos-aquae* (AnaS). The geometry, adapted from the cryo-EM images, comprises a cylindrical shell with conical ends, with height and diameter, respectively, of 500 nm and 85 nm. The protein wall was idealized as a continuum shell with a thickness of 2.8 nm and a shell density of 1350 kg/m^3^. The rib-like structure of the gas vesicle wall was mirrored in the computational model by an elastic anisotropic material model, with elastic moduli across and along the principal axis of the GV of 0.98 GPa and 3.92 GPa, respectively (Maresca et al. 2017). In order to simulate the nearly incompressible nature of the protein shell, we assigned a Poisson’s ratio of 0.499. We note that the material parameters were not obtained from direct experimental measurements, but rather chosen such that, in addition to falling within a range of parameters consistent with those of protein-based biological materials (Gosline et al. 2002), they effectively replicated the buckling pressures observed experimentally.

We next added a helical rod that spirals around the cylindrical portion of the GV shell, modeling the GvpC molecules. We modeled the GvpC rod as a shell of radius 0.6 nm. The helical structure was generated by assigning a pitch of 4.9 nm. The finite element model of the resultant wild type GV was obtained by discretizing the entire geometry with quadrilateral shell elements of effective side length 1 nm with reduced integration (*i.e*., S4R elements) in Abaqus (Dassault Systemes Simulia, France). These general-purpose shell elements with only one integration point within each element are capable of capturing both tensile and in-plane bending, and, with a sufficiently fine mesh size, are computationally cost-effective. We subjected the interior surfaces of the GV to an initial pressure of 101 kPa, modeling the inner gas pressure. We further subjected the vertices at both the top and bottom conical ends of the GV to a zero-displacement Dirichlet boundary condition, which prevented rigid body translations and rotations of the entire GV structure.

In order to investigate the effect of GvpC density on the buckling pressure, we first computed the total length of the helix where N, D, and z are the total number of turns, the perimeter of the GV cross-section, and the pitch of the helix, respectively. Given the pitch and the length of the cylindrical segment of the GV model, 416.5 nm, the total number of turns was computed as 85. We thus computed the total length of the helix as 22.702 micrometers. Given that the length of GvpC is ∼25 nm, about 908 GvpC molecules constituted the helix in our model. We generated six additional finite element models with distinct GvpC saturation levels of 90%, 80%, 60%, 40%, 20%, and 10%, for which we randomly removed about 90, 180, 360, 540, 720, and 810 GvpC units, respectively.

We conducted linear buckling analysis (LBA) and solved the corresponding eigenvalue problem to obtain the threshold buckling pressures for each model. We solved this problem using the Lanczos algorithm and obtained the first ten modes of buckling. Unlike the buckling modes (*i.e*., eigenvectors), which were virtually identical at different levels of GvpC saturation, the buckling pressures (*i.e*., eigenvalues) were remarkably dependent on the GvpC density, with an almost linear monotonic relation, where decreasing the saturation level decreases the buckling pressure. Figure S11 depicts the buckling modes and pressures for 100%, 60%, 20%, and 0% GvpC saturations.

### Bioinformatics and visualization

Protein sequence alignment was carried out using Clustal Omega (Sievers et al. 2011) and visualized with Jalview (Waterhouse et al. 2009). Protein conservation analysis was performed using ConSurf (Ashkenazy et al. 2016). Data were visualized using GraphPad Prism, IMOD (Kremer, Mastronarde, and McIntosh 1996), Chimera (Pettersen et al. 2004), and ChimeraX (Goddard et al. 2018). Identified crosslinks were visualized using xiNET(Combe, Fischer, and Rappsilber 2015).

## ACKNOWLEDGMENTS

The authors are grateful to Catherine Oikonomou for helpful editorial comments. We thank Songye Chen for assistance with tomography data collection. Electron microscopy was performed in the Beckman Institute Resource Center for Transmission Electron Microscopy at Caltech. The Proteome Exploration Laboratory (PEL) is supported by the Beckman Institute and NIH 1S10OD02001301. This work was supported by the National Institutes of Health (grant R01-AI127401 to G.J.J. and R01-EB018975 to M.G.S.) and the Caltech Center for Environmental Microbial Interactions (CEMI). Related research in the Shapiro Laboratory is supported by the Packard Foundation, the Chan Zuckerberg Initiative and the Heritage Medical Research Institute.

## AUTHOR CONTRIBUTIONS

P.D. conceived experiments, prepared samples, acquired and analyzed data, performed data exploration, drafted the manuscript, and prepared the figures. L.A.M initiated the project and collected data for Mega GVs. R.C.H performed mutation screening for GvpA and participated in initial sample preparation and optimization for Mega GVs. H.S. performed finite element simulation and analyzed data. T-Y.W performed XLMS experiments and analyzed the data.

D.M expressed and purified GV samples. G.L. participated in initial sample preparation and optimization for Mega GVs. T-F.C. supervised XLMS experiments. All authors participated in correction of the manuscript. M.G.S. participated in guidance, experimental design, funding, and correction/advising on writing the manuscript. G.J.J participated in guidance, experimental design, funding, and correction/advising on writing the manuscript.

## DATA AVAILABILITY

Cryo-ET density maps and GvpA/GvpC integrative model are available on Zenodo zenodo.org/record/6820642#.Ys0aROzMKw1.

## SUPPLEMENTARY MATERIALS

**Figure S1.**
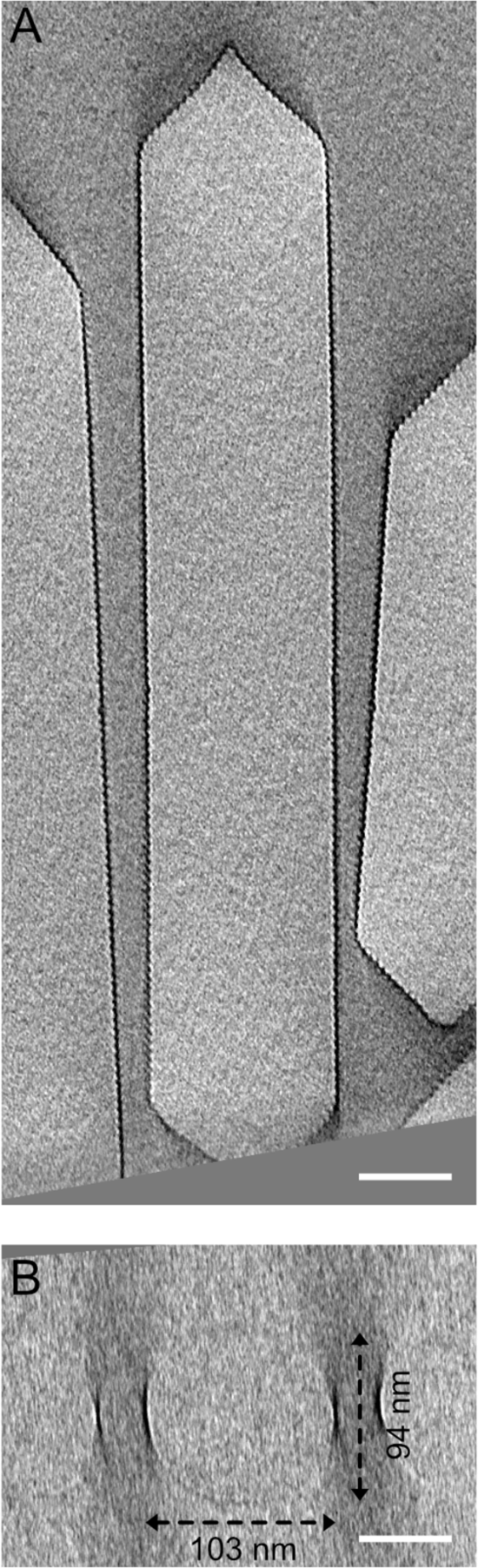
GV flattening in the thin ice. (**A**) XY and (**B**) XZ tomographic slices of the deformed Ana GV. Scale bars, 50 nm.

**Figure S2.**
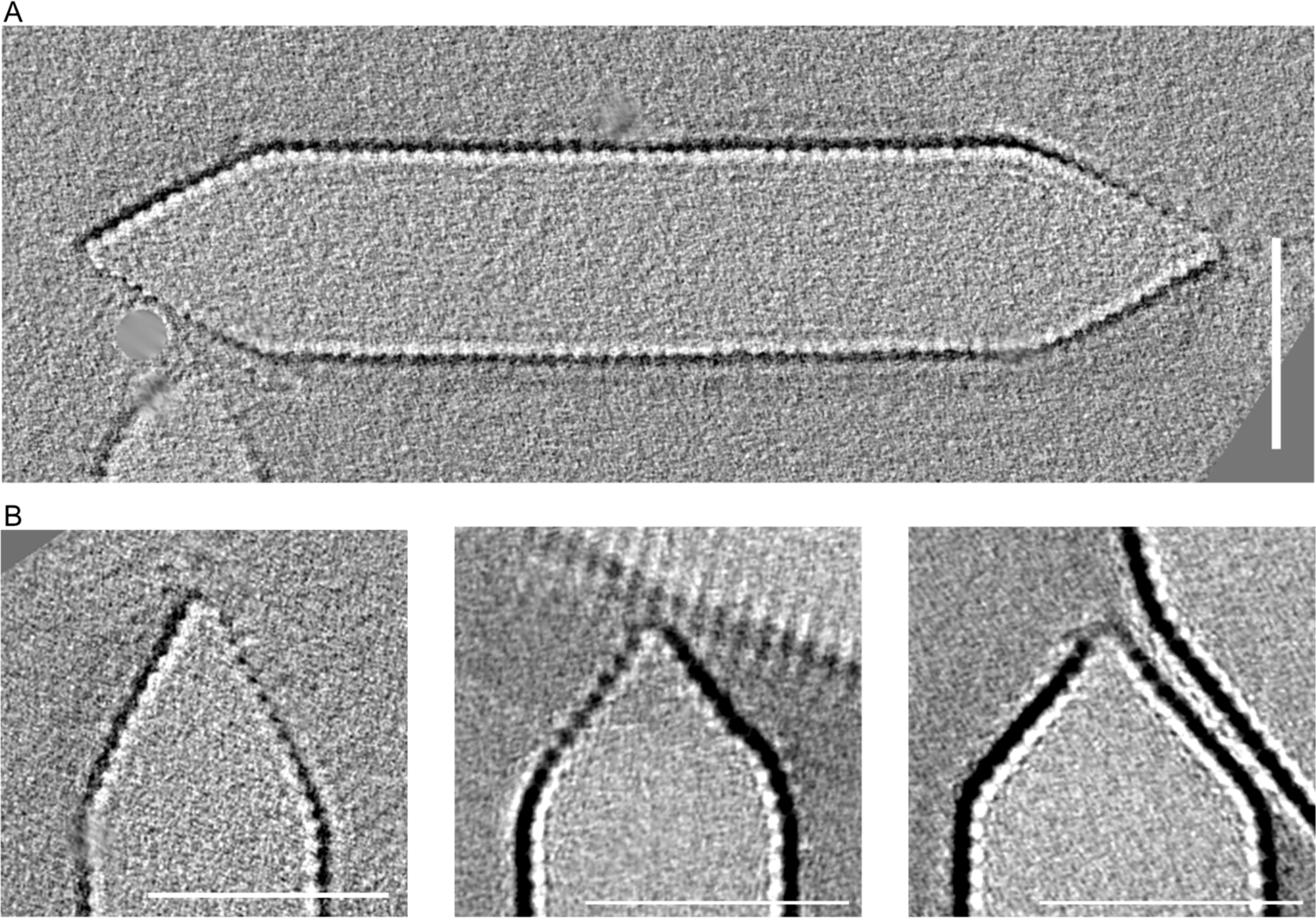
The architecture of Mega GVs. (**A**) Representative central slice from cryo-electron tomogram of individual Mega GV. (**B**) Central tomographic slices of the Mega GV conical ends with slightly different shapes. Scale bars, 50 nm.

**Figure S3.**
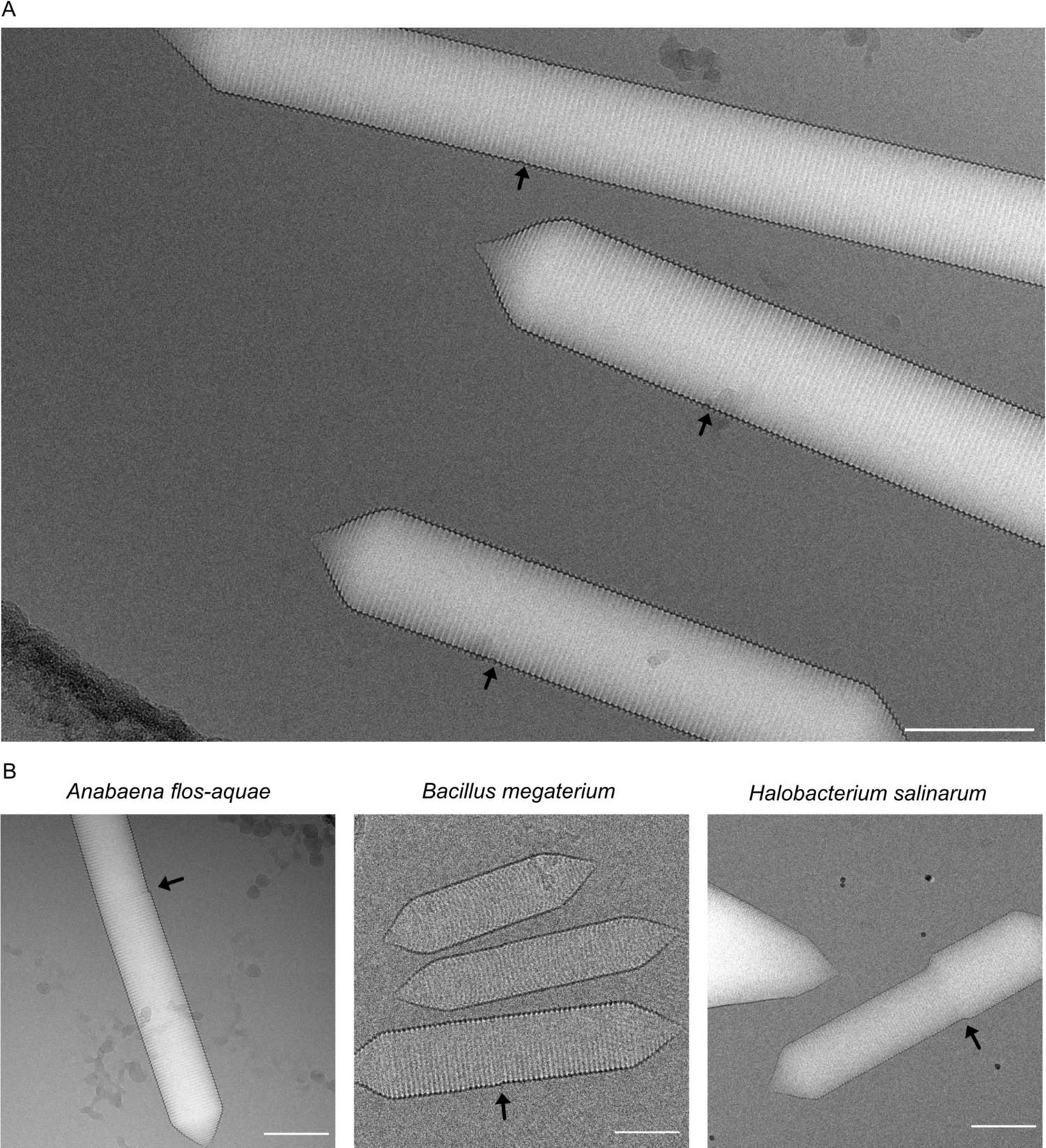
GV Polarity inversion point. (**A**) Location of the polarity inversion point. (**B**) GVs can have different diameters on either side of the inversion point. The black arrows indicate the location of the inversion point.

**Figure S4.**
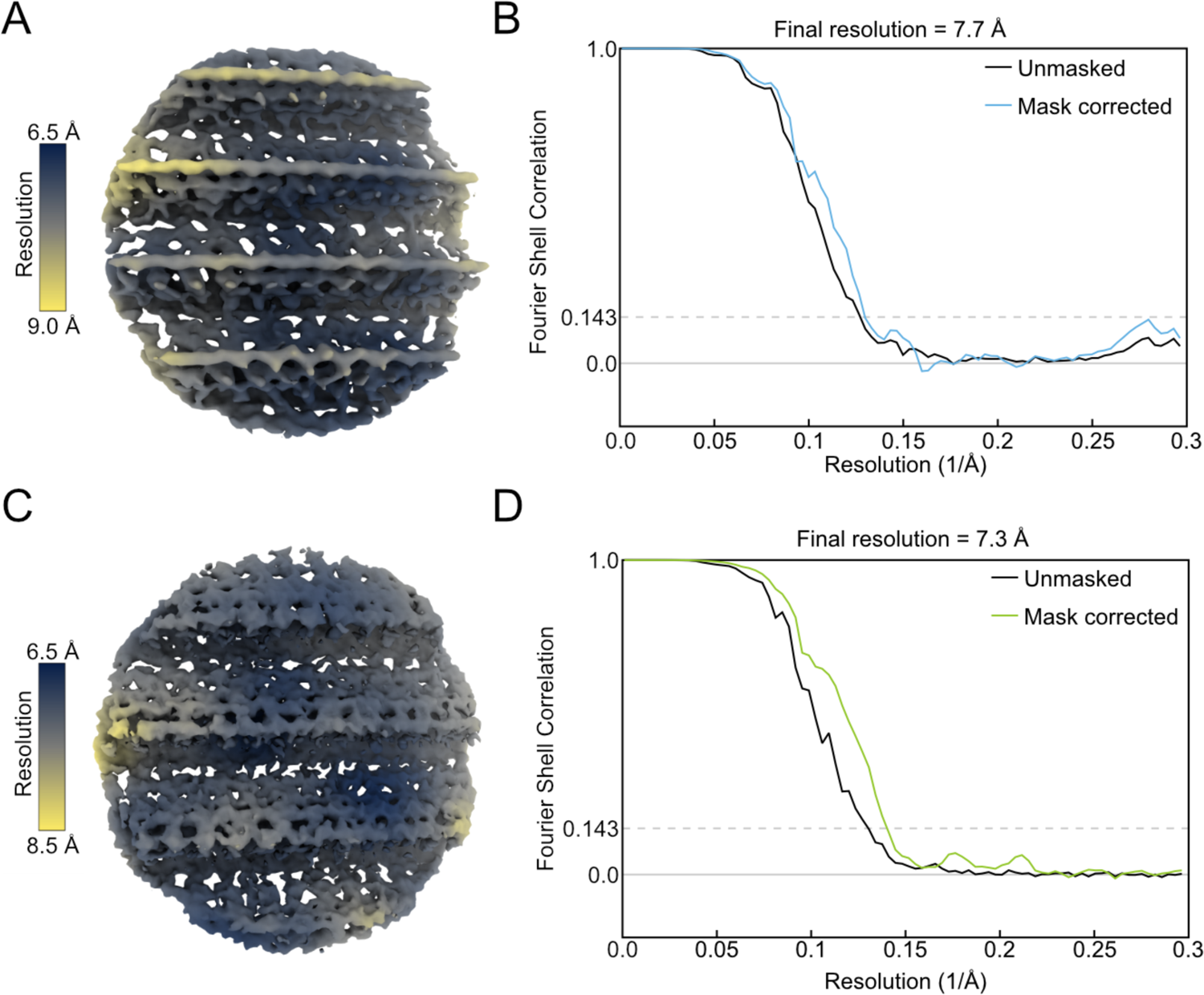
Resolution estimation of the subtomogram of Ana and AnaS GV shells. (**A**,**C**) Local resolution estimation of subtomogram averages for Ana (A) and AnaS (C) GV shell. (**B, D**) Global FSC for Ana (B) and AnaS (D) subtomogram averages.

**Figure S5.**
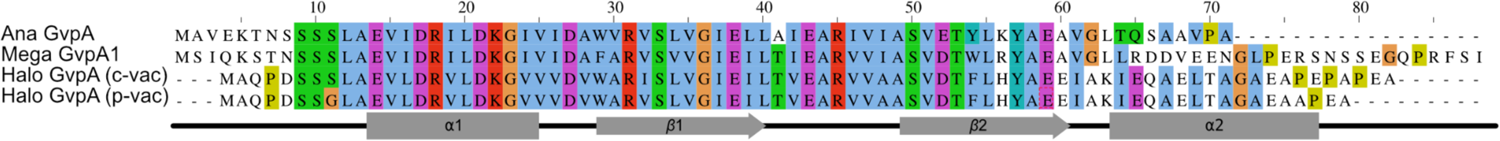
Protein sequence alignment. Sequence alignment among homologs of the major structural protein (GvpA) from Mega, Ana, and Halo (p-vac and c-vac gene clusters).

**Figure S6.**
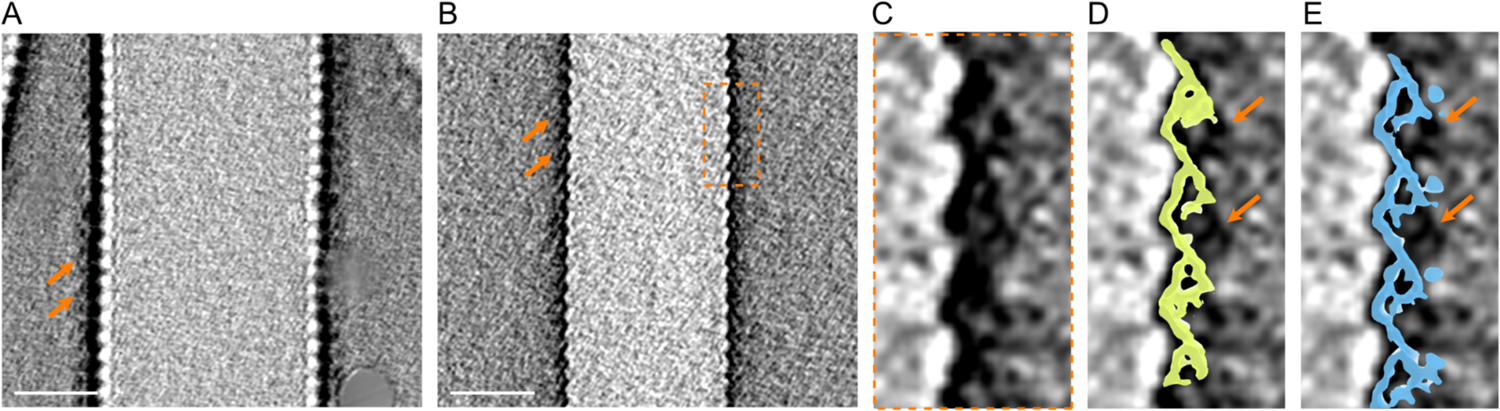
Additional densities on the surface of Mega GVs. (**A**,**B**) Slices from cryo-electron tomograms of individual Mega GVs show additional density on the surface. Defocus values: (**A**) −5 µm and (**B**) −1 µm. (**C**) Enlarged section form **B** as outlined by orange dashed box. (**D, E**) Superimposition of subtomogram averages (Figure 3D) for AnaS and Ana GV shell. Orange arrows indicate extra densities. Scale bars, 20 nm.

**Figure S7.**
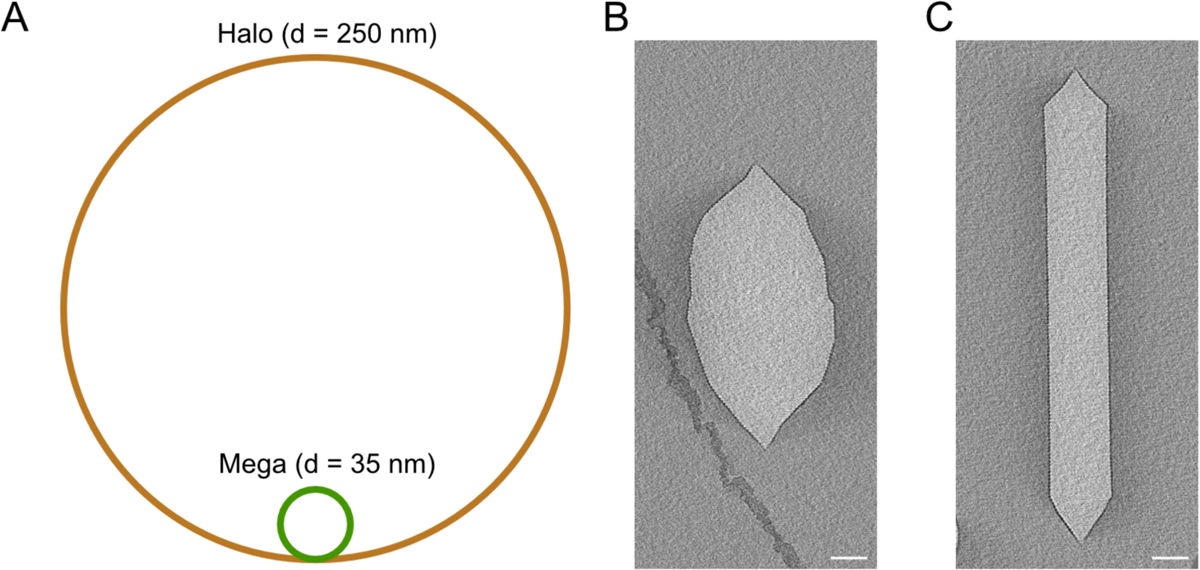
GVs adopt a wide range of diameters and different morphologies. (**A**) Schematic showing difference in rib curvature between smallest (Mega) and largest (Halo) measured diameter (Dutka et al. 2021). (**B**,**C**) Representative central slices from cryo-electron tomograms of individual Halo GVs encoded by (**B**) c-vac and (**C**) p-vac gene clusters. Scale bars, 50 nm.

**Figure S8.**
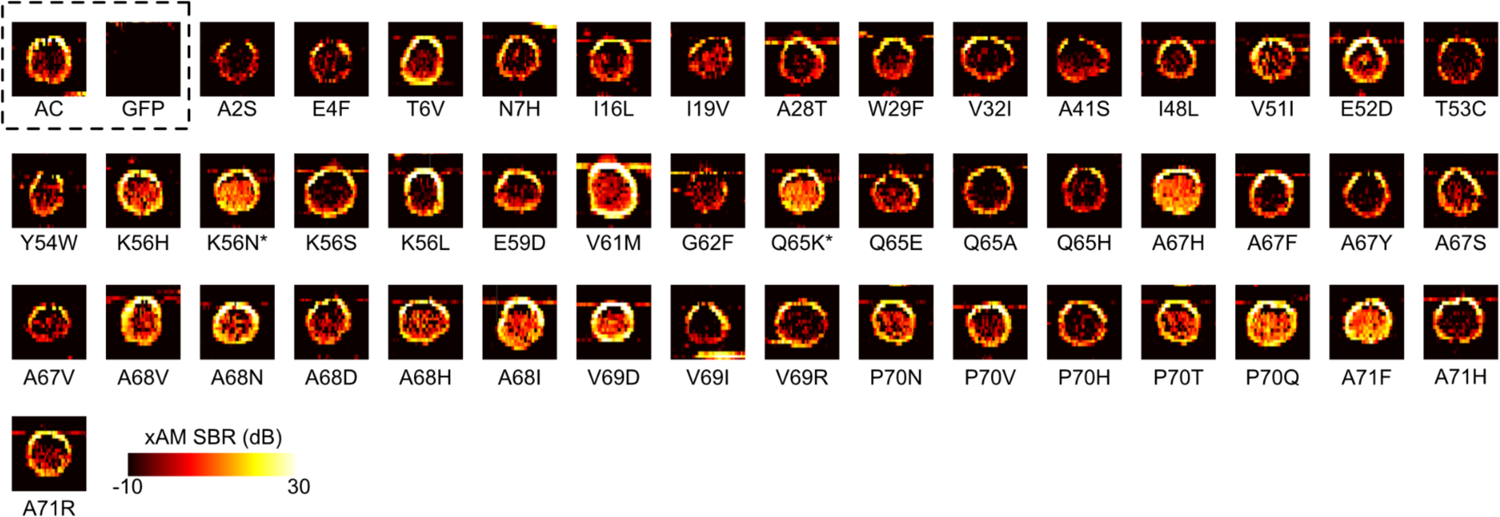
Ultrasound images of *E. coli* clones expressing select GvpA mutants. Pre-minus-post-collapse nonlinear xAM images of clones of *E. coli* expressing GVs with the indicated mutations in GvpA. All the shown mutants display clear non-zero contrast and therefore successfully form GVs. Wild type GvpA (AC) and GFP are included as positive and negative controls, respectively. Color map corresponds to SBR, the signal-to-background ratio. *Mutations K56N and Q65K occurred in the same clone. GV expression is more pronounced on the edges of the patches because of those cells’ increased access to nutrients.

**Figure S9.**
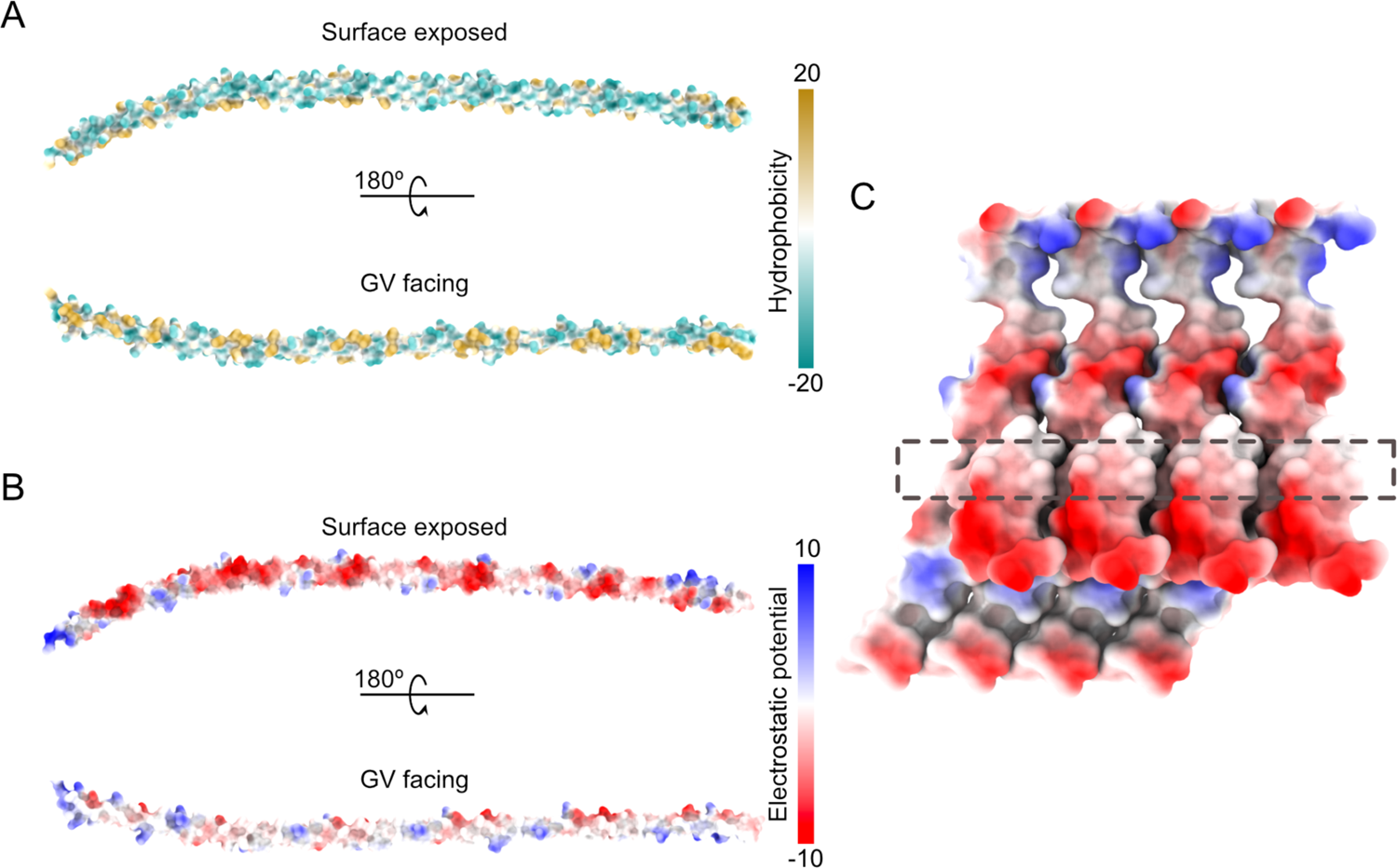
Hydrophobicity and charge distribution on GvpC surface. (**A**,**B**) AlphaFold2 predicted models of full-length GvpC. (**A**) Hydrophobicity of the GvpC surface. (**B**) Distribution of the electrostatic potential of the GvpC surface. (**C**) Distribution of the electrostatic potential on the GvpC binding model.

**Figure S10.**
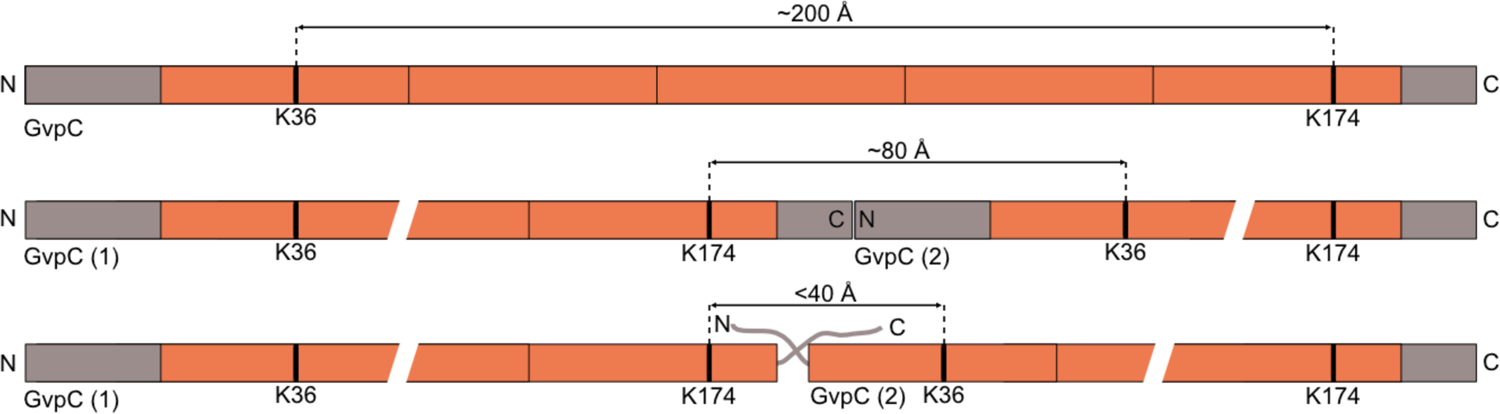
Distances for different scenarios of Lys cross-linking between GvpC molecules.

**Figure S11.**
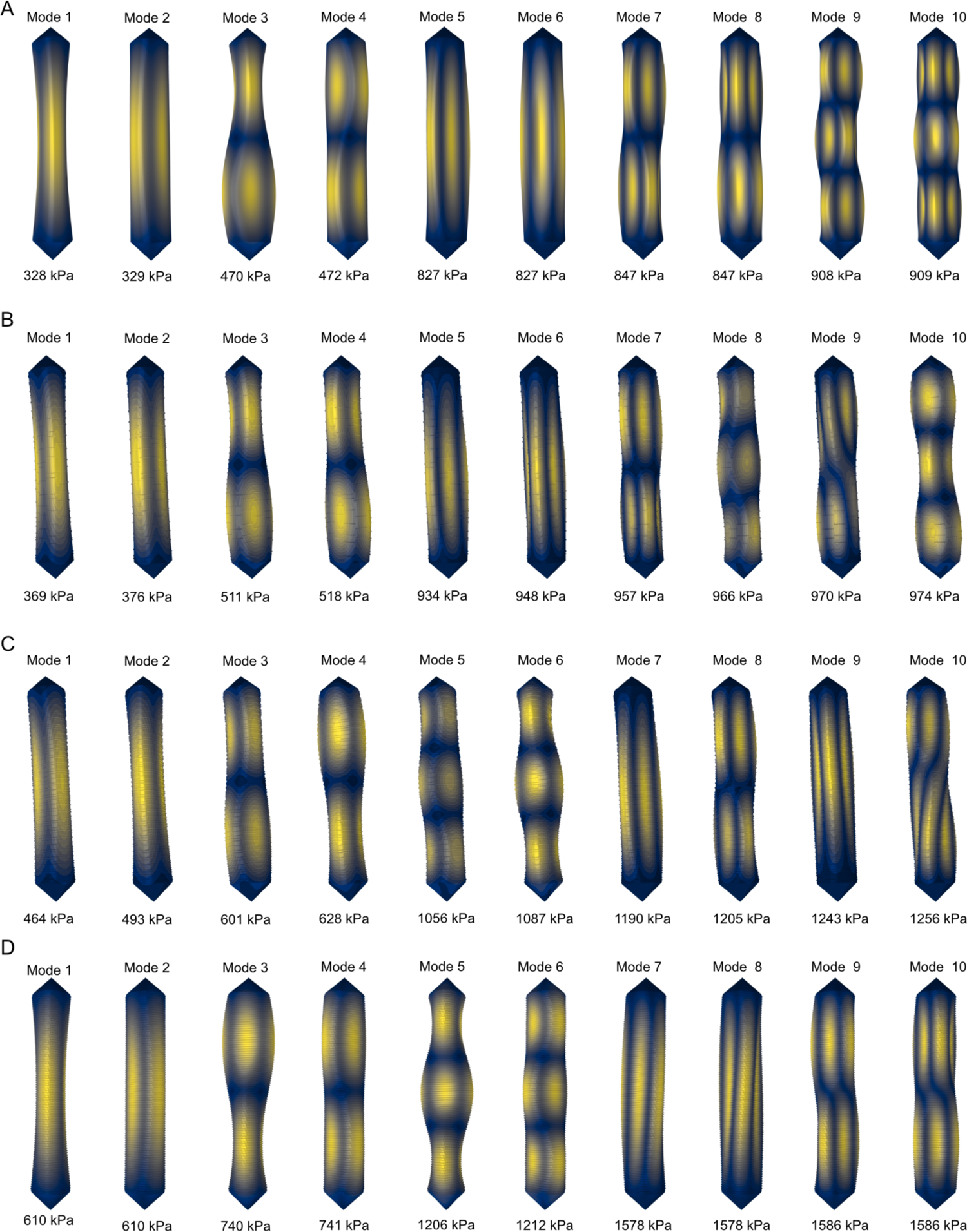
Buckling modes of GVs with different degrees of GvpC saturation. The first ten buckling modes and pressures were obtained from linear buckling analysis for GV with distinct saturation levels of GvpC. Rows from top to bottom represent GvpC densities of (**A**) 0%, (**B**) 20%, (**C**) 60%, and (**D**) 100%.

**Table S1.** Data collection and processing parameters for GV shell.

|  | Native GVs | AnaS GVs |
| --- | --- | --- |
| Magnification | 53,000× | 53,000× |
| Voltage (keV) | 300 | 300 |
| Energy Filter | Yes | Yes |
| Slit width (eV) | 20 | 20 |
| Pixel size (Å) | 0.8435 | 0.8435 |
| Defocus range (μm) | 1.5 to 3.5 | 1.5 to 3.5 |
| Defocus step (μm) | 0.5 | 0.2 |
| Tilt range (min/max, step) | -60/60°, 3° or -44/44°, 4° | -45/45°, 3° |
| Tilt scheme | Dose-symmetric | Dose-symmetric |
| Total dose (electrons/Å<sup>2</sup>) | ~45 | ~45 |
| Frame number | 10 or 5 | 10 |
| Tomograms used/acquired | 81/368 | 28/103 |
| Number of cylinders | 136 | 32 |
| Final subtomograms (no.) | 73,878 | 27,184 |
| Symmetry | C1 | C1 |
| Map resolutions<br>(FSC = 0.143) | 7.7 | 7.3 |

**Table S2.**
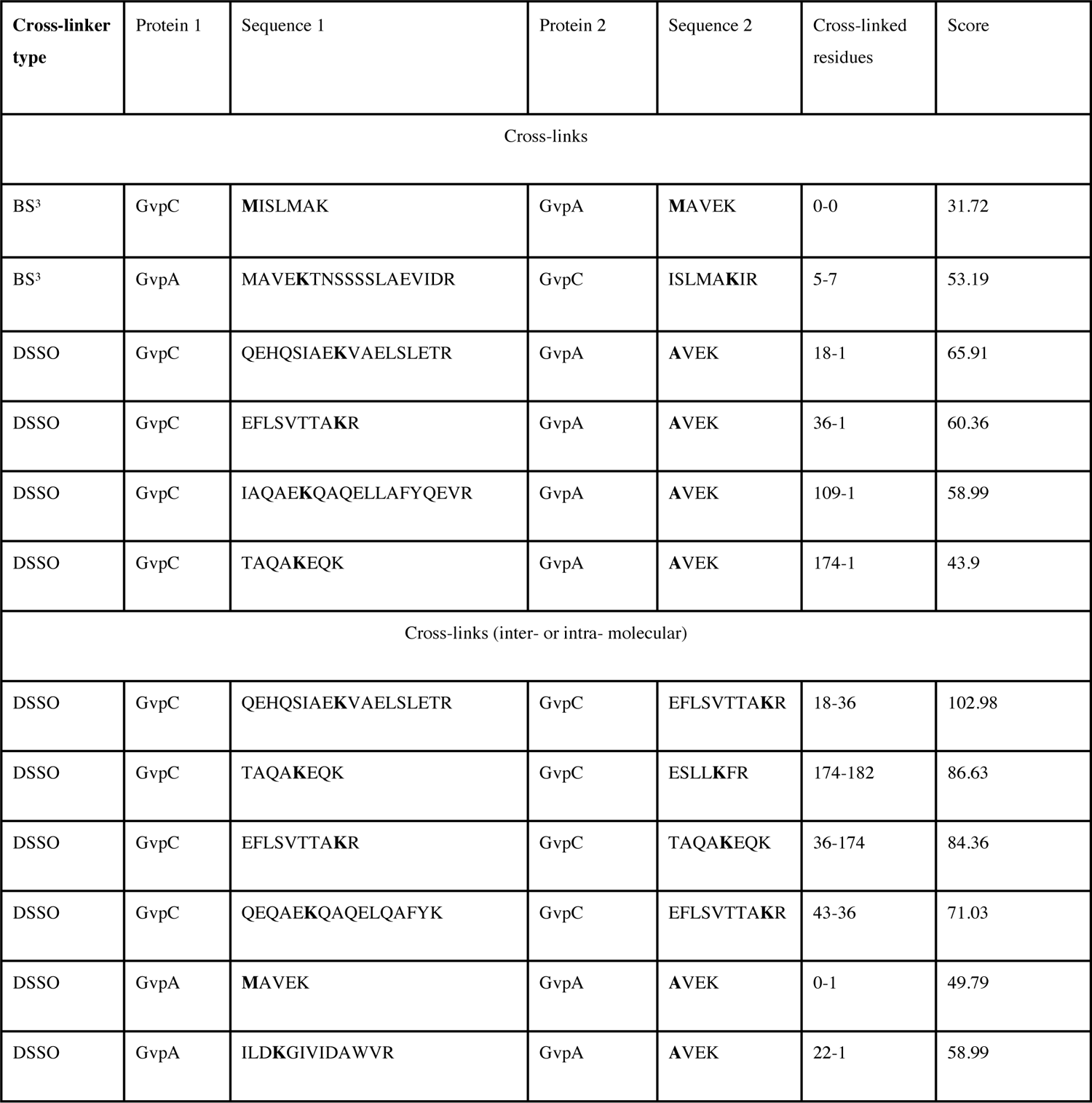
List of validated cross-linked peptides.

